# Polyamines mediate enterovirus attachment directly and indirectly through cellular heparan sulfate synthesis

**DOI:** 10.1101/2021.10.31.466121

**Authors:** Bridget M. Hulsebosch, Oreoluwa S. Omoba, Natalie J. LoMascolo, Bryan C. Mounce

**Author notes:** Corresponding author: Department of Microbiology and Immunology, Loyola University Chicago, Stritch School of Medicine, 2160 S. First Ave., Maywood, IL 60153.

## Abstract

Productive viral infection begins with attachment to a susceptible cell, and viruses have evolved complex mechanisms to attach to and subsequently enter cells. Prior to engagement with a cellular receptor, viruses frequently interact with nonspecific attachment factors that can facilitate virus-receptor interactions and viral entry. Polyamines, small positively-charged molecules abundant in mammalian cells, mediate viral attachment, though the mechanism was not fully understood. Using the Coxsackievirus B3 (CVB3) enterovirus model system, we show that polyamines mediate viral attachment both directly and indirectly. The polyamine putrescine specifically enhances viral attachment to cells depleted of polyamines. Putrescine’s positive charge mediates its ability to enhance viral attachment, and polyamine analogs are less efficient at mediating viral attachment. In addition to this direct role of polyamines in attachment, polyamines facilitate the cellular expression of heparan sulfates, negatively-charged molecules found on the cell surface. In polyamine-depleted cells, heparan sulfates are depleted from the surface of cells, resulting in reduced viral attachment. We find that this is due to polyamines’ role in the process of hypusination of eukaryotic initiation factor 5A, which facilitates cellular translation. These data highlight the important role of polyamines in mediating cellular attachment, as well as their function in facilitating cellular heparan sulfate synthesis.

## Introduction

Viral attachment to a susceptible cell is the first step in the complex process of infection. Viruses have distinct mechanisms to attach to cells, evolving affinities for cellular attachment factors and receptors. In the case of enteroviruses, several ubiquitous molecules serve as nonspecific attachment factors, including heparan sulfates^1–6^, vimentin^7,8^, and sialic acids^9,10^. These molecules serve to enhance virus-cell interaction prior to receptor engagement. The initial interaction with these nonspecific molecules serves to convert a three-dimensional search for the viral receptor into a two-dimensional search, enhancing the potential for engagement with the specific viral receptor. Coxsackievirus, one such enterovirus, engages with these cell surface molecules prior to entry mediated by a receptor, the Coxsackievirus and adenovirus receptor (CAR), to initiate infection^4,10^. Coxsackieviruses also interact with decay accelerating factor (DAF, CD55), which also mediates Coxsackievirus attachment and entry^11–13^.

A common childhood infectious agent, Coxsackievirus infection frequently resolves without the need for intervention. However, CVB3 causes significant disease, including hand foot and mouth disease, meningitis, encephalitis, conjunctivitis, and myocarditis^14–16^. Coxsackievirus’ ability to infect and persist in cardiac tissue represents not only a threat to children but also adults^17,18^. Seroprevalence is high in some areas, reaching levels as high as 50%^19^. Frequent outbreaks of enteroviruses such as enterovirus-A71 or -D68 highlight the ability of these viruses to rapidly spread and cause significant morbidity and mortality. Unfortunately no antivirals or vaccines are available to treat or prevent infection.

We previously showed that Coxsackievirus B3 (CVB3) attachment to cells requires polyamines^20^, small aliphatic molecules, comprised of short carbon chains and tertiary amines. Polyamines function in cell cycling, translation, and nucleotide metabolism within cells^21^, and they’re also important for CVB3 infection^22^. In polyamine-depleted conditions, CVB3 replication is significantly attenuated, both *in vitro* and using the *in vivo* mouse model^22^. By passaging CVB3 in polyamine-depleted cells, we previously observed four escape mutants, three in the viral protease 2A or 3C^23,24^, and one in the capsid protein VP3^20^, suggesting a role in viral attachment and/or entry. Further examination revealed a global role for polyamines in attachment of diverse enteroviruses; however, the mechanism remains to be fully understood.

Here, we show that CVB3 relies on polyamines to mediate viral attachment both directly and indirectly. CVB3 attachment to polyamine depleted cells is diminished using several inhibitors of polyamine metabolism. However, incubation of virus with polyamines enhances viral attachment in a dose-dependent manner. Using a panel of natural and synthetic polyamines, we find that specifically the natural polyamine putrescine enhances viral attachment, though spermidine and spermine also function to a slightly reduced degree. We find that the viral escape mutant CVB3 VP3^Q234R^, which is resistant to polyamine depletion, does not require polyamines, nor do polyamines enhance attachment. However, CVB3 with a negative or neutral charge at this amino acid do rely on putrescine. Finally, we find that cellular factors also contribute to viral attachment. Heparan sulfate, a nonspecific attachment factor, is reduced in cell surface abundance in polyamine-depleted cells, which likely depends on polyamines for the translation of a heparan sulfate synthetic enzyme. Together, these results demonstrate roles for polyamines in directly and indirectly mediating CVB3 attachment, highlighting the importance of these molecules in CVB3 infection.

## Results

### Polyamines facilitate CVB3 attachment

We previously demonstrated that polyamines facilitate viral attachment^20^, though the mechanism was not clear. To recapitulate these results, we performed a viral attachment assay by applying virus to cells on ice for a five-minute period before washing away unbound virus. Immediately after washing, cells were overlaid with agarose-containing media to limit virus spread. To this attachment assay, we added increasing doses of DFMO to deplete polyamines, and we observed a significant reduction in attached virus with increasing DFMO concentrations (Figure 1A, representative plaques shown in Figure 1B). Importantly, in this assay, polyamines are replenished in the agarose-containing media, so polyamine depletion is limited strictly to the binding phase of infection.

**Figure 1.**
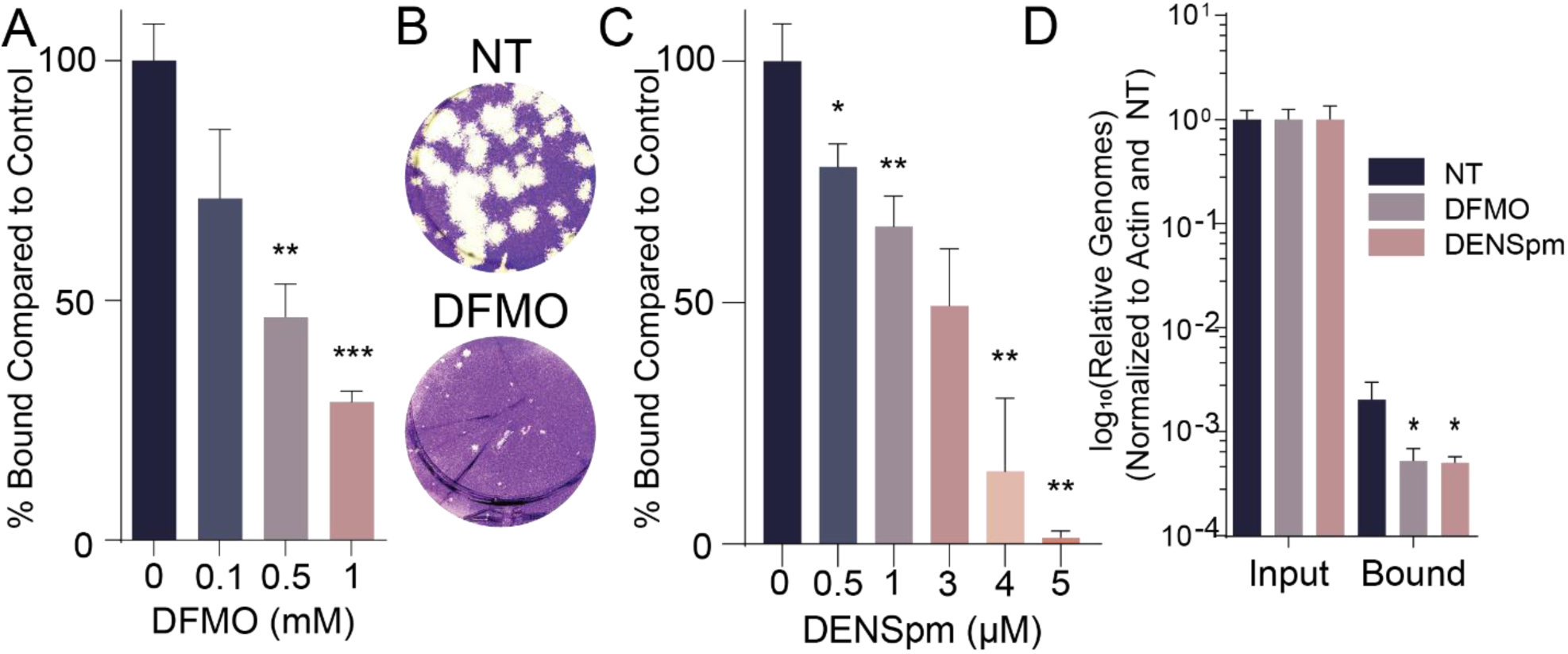
Polyamines facilitate CVB3 attachment. (A) Vero cells were treated with increasing doses of DFMO for four days prior to measuring viral attachment by plaque assay. Bound virus was quantified and compared to untreated conditions. (B) Representative image of assay as performed in (A). (C) Vero cells were treated with increasing doses of DENSpm 16h prior to performing a binding assay as in (A). (D) Cells were treated with DFMO or DENSpm, infected with CVB3, and bound virus quantified by qRT-PCR after washing away unbound virus. *p<0.05, **p<0.01, ***p<0.001 by Student’s T test (N≥3).

While DFMO treatment depletes polyamines by inhibiting ODC1, polyamines can also be depleted by the drug diethylnorspermidine (DENSpm), which activates polyamine catabolism through the enzyme spermidine-spermine acetyltransferase. To determine if DENSpm similarly restricted virus attachment, we treated cells with increasing doses of DENSpm and measured virus attachment as with DFMO. Again, we observed a dose-dependent reduction in virus attachment (Figure 1C, representative plaques in Figure 1D), suggesting that polyamine depletion, and not DFMO or DENSpm themselves, reduces viral attachment. To confirm these results with a more specific attachment assay, we performed the assay as previously, treating with both DFMO and DENSpm, but immediately after virus attachment and washing of excess virus, we collected cells and bound virus in Trizol, purified RNA, reverse transcribed, and measured viral genomes via qRT-PCR, normalizing to cellular actin. Again, we observed a significant reduction in virus attachment in polyamine depleted cells, both for DFMO and DENSpm treatment (Figure 1E), again implicating polyamines in viral attachment.

### Exogenous polyamines rescue virus replication and attachment in DFMO-treated cells

Cells acquire polyamines either through synthesis or via dedicated transporters on the cellular surface. We previously demonstrated that applying exogenous polyamines to cells rescue virus replication, including for CVB3. To determine if specific polyamines enhance virus replication, possibly indicating specificity in polyamine-virus interactions, we added individual polyamines to DFMO-treated cells and measured viral titers after a 24-h infection. When we treated cells with any of the biogenic polyamines (putrescine, spermidine, spermine, or a mix of the three), we observed a full rescue in virus replication (Figure 2A). Interestingly, when we applied cadaverine, norspermidine, or propylamine, we observed a modest rescue of virus titers. These data suggest that CVB3 replication relies on the three polyamines synthesized by eukaryotic cells.

**Figure 2.**
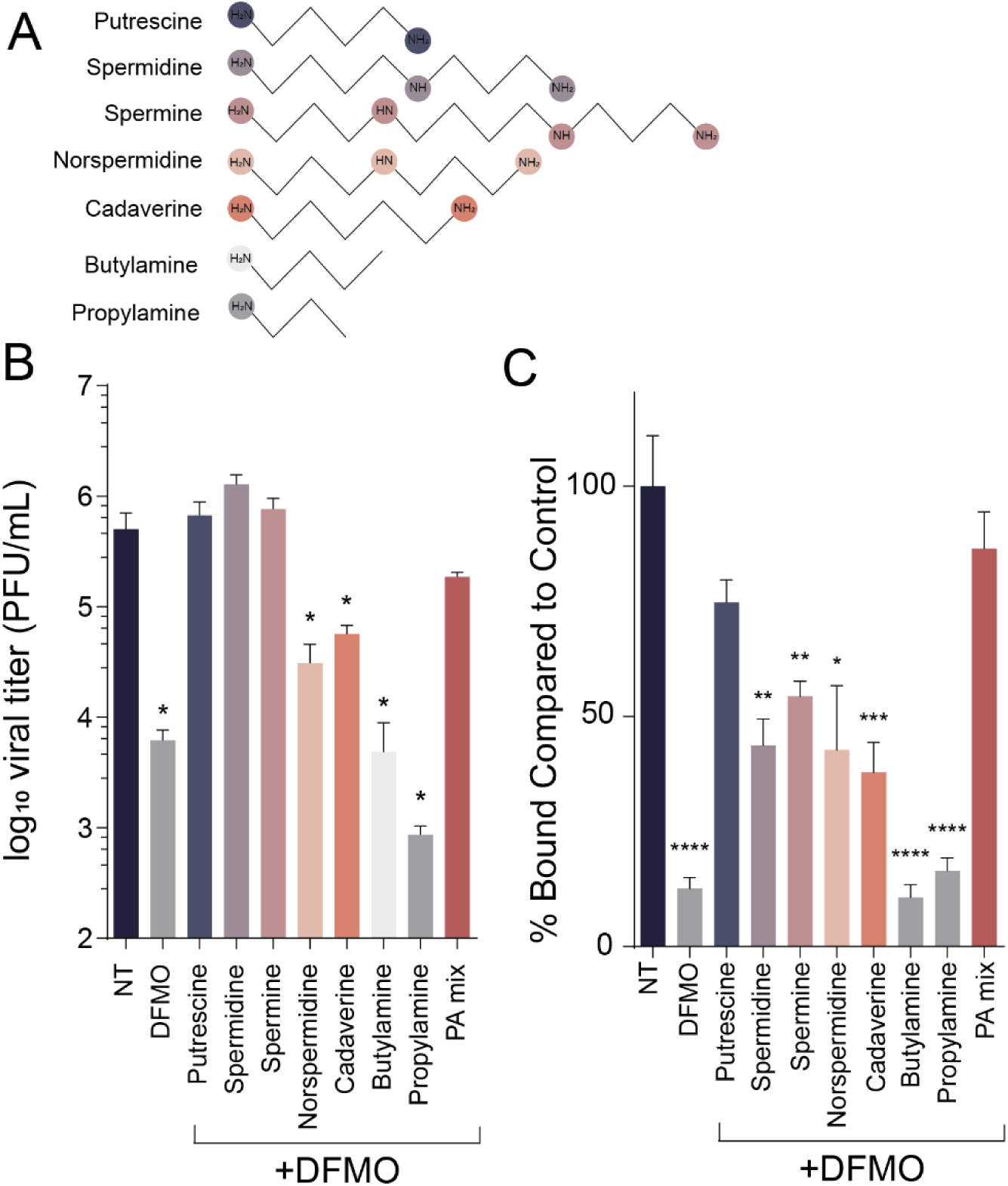
Exogenous polyamines rescue CVB3 replication and attachment in DFMO-treated cells. Vero cells were treated with DFMO for four days prior to supplementation with exogenous polyamines as shown in (A). Cells were subsequently infected and (B) viral titers were measured 24 hpi. (C) Viral attachment was measured in cells treated as in (A). *p<0.05, **p<0.01, ***p<0.001 by Student’s T test (N≥2).

To more precisely determine which polyamine, if any in particular, enhanced virus attachment, we performed an attachment assay with the individual polyamines, adding the polyamines to the viral inoculum prior to applying to the cells. When we did this, we observed that curiously only putrescine rescued virus attachment (Figure 2B, representative plaques shown in Figure 2C). Additionally, when we mixed the three eukaryotic polyamines (including putrescine), we observed a full rescue for attachment. These data suggest that specifically putrescine enhances CVB3 attachment.

### Putrescine enhances CVB3 binding

Having observed that specifically putrescine enhances virus attachment, we next investigated whether incubating polyamines directly with virus prior to attachment mediated this rescue. To this end, we incubated CVB3 with a panel of polyamines prior to attachment to cells on ice for five minutes, in our standard attachment assay. We observed that a mixture of the biogenic polyamines (“PA Mix”) enhanced attachment, but only putrescine enhanced attachment, while none of the other polyamines were functional (Figure 3A). To determine if this was concentration dependent, we incubated CVB3 with increasing doses of putrescine as before. When we did this, we observed a dose-dependent rescue of virus attachment in DFMO-treated cells. In a similar vein, we incubated CVB3 with increasing doses of spermidine and spermine, and we observed a modest rescue of virus attachment (Figure 3C, D). Together, these data suggest that specifically putrescine mediates viral attachment when directly incubated with CVB3.

**Figure 3.**
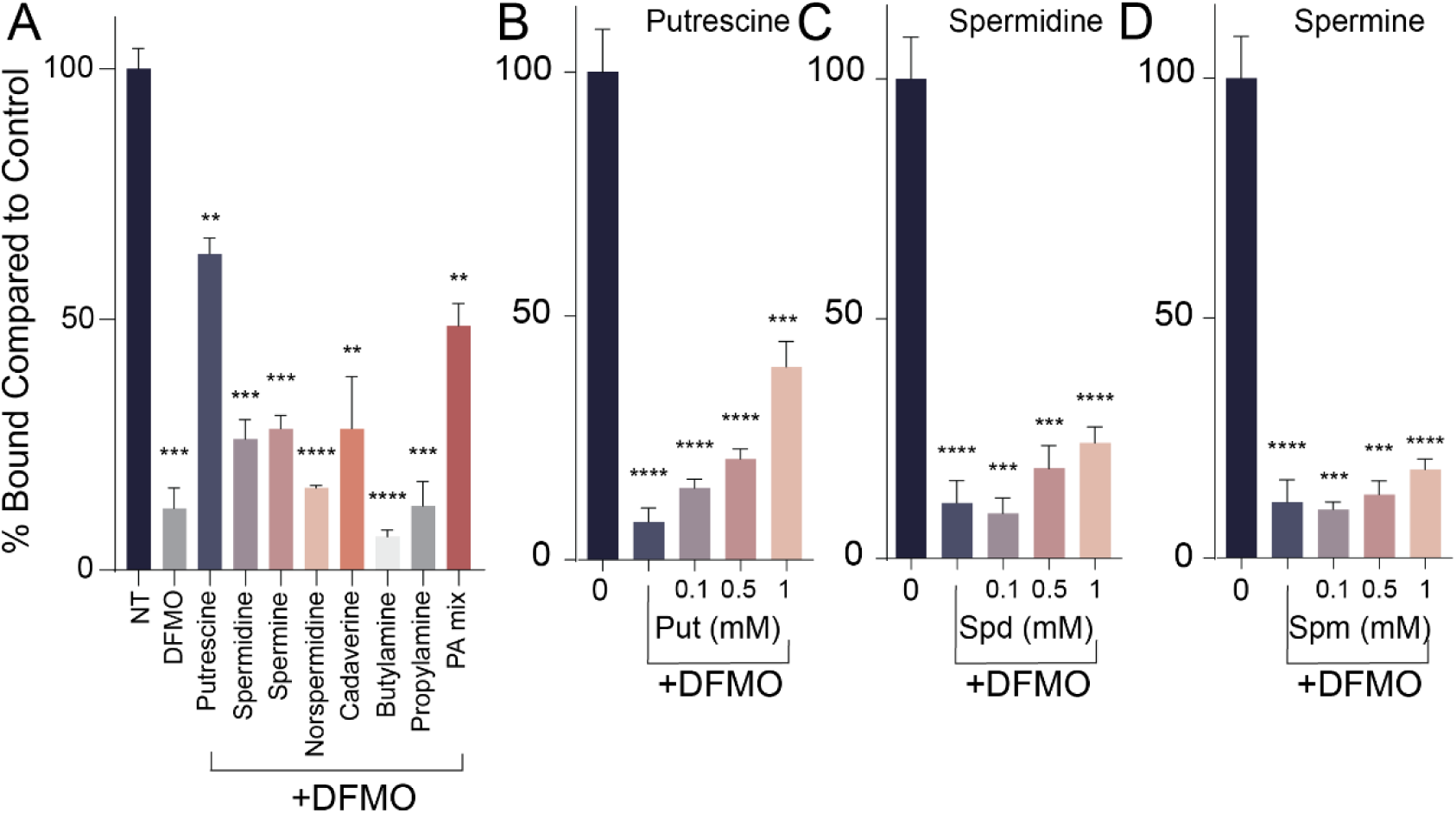
The biogenic polyamines facilitate CVB3 attachment in a dose-dependent manner. (A) Vero cells were treated with 1 mM DFMO for four days prior to infection with CVB3 that was incubated with 5 mM of the indicated polyamines. Viral attachment assays were subsequently performed. (B-D) cells were treated as in (A) but virus was incubated with increasing doses of (B) putrescine, (C) spermidine, and (D) spermine. Unattached virus was washed and attached virus revealed and quantified after plaque development. *p<0.05, **p<0.01, ***p<0.001, ****p<0.0001 by Student’s T test (N≥3).

### VP3^Q234R^ mediates attachment independently of polyamines

We previously identified a CVB3 mutant that attached to cells independently of polyamines. The mutant in VP3 at glutamine 234 altered the negatively-charged glutamine to a positively-charged arginine (hereafter referred to as VP3^Q234R^). We reasoned that polyamine depletion limits positively-charged polyamines within the cell and that the virus responds to the polyamine depletion through the incorporation of this positively-charged amino acid. Interestingly, we observed this phenotype in other mutants resistant to polyamine depletion, both for CVB3 and CHIKV. To determine if this mutant attached to cells independently of polyamines, we incubated CVB3 VP3^Q234R^ with increasing doses of polyamines and performed an attachment assay. When we did this, we observed that the amount of attached virus did not change with putrescine concentration (Figure 4A), suggesting that this mutant does not rely on polyamines, putatively because of this positively-charged amino acid. In contrast, when we performed these assays with VP3^Q234A^ or VP3^Q234E^ mutants, we observed continued dependence on putrescine for attachment (Figure 4B, C).

**Figure 4.**
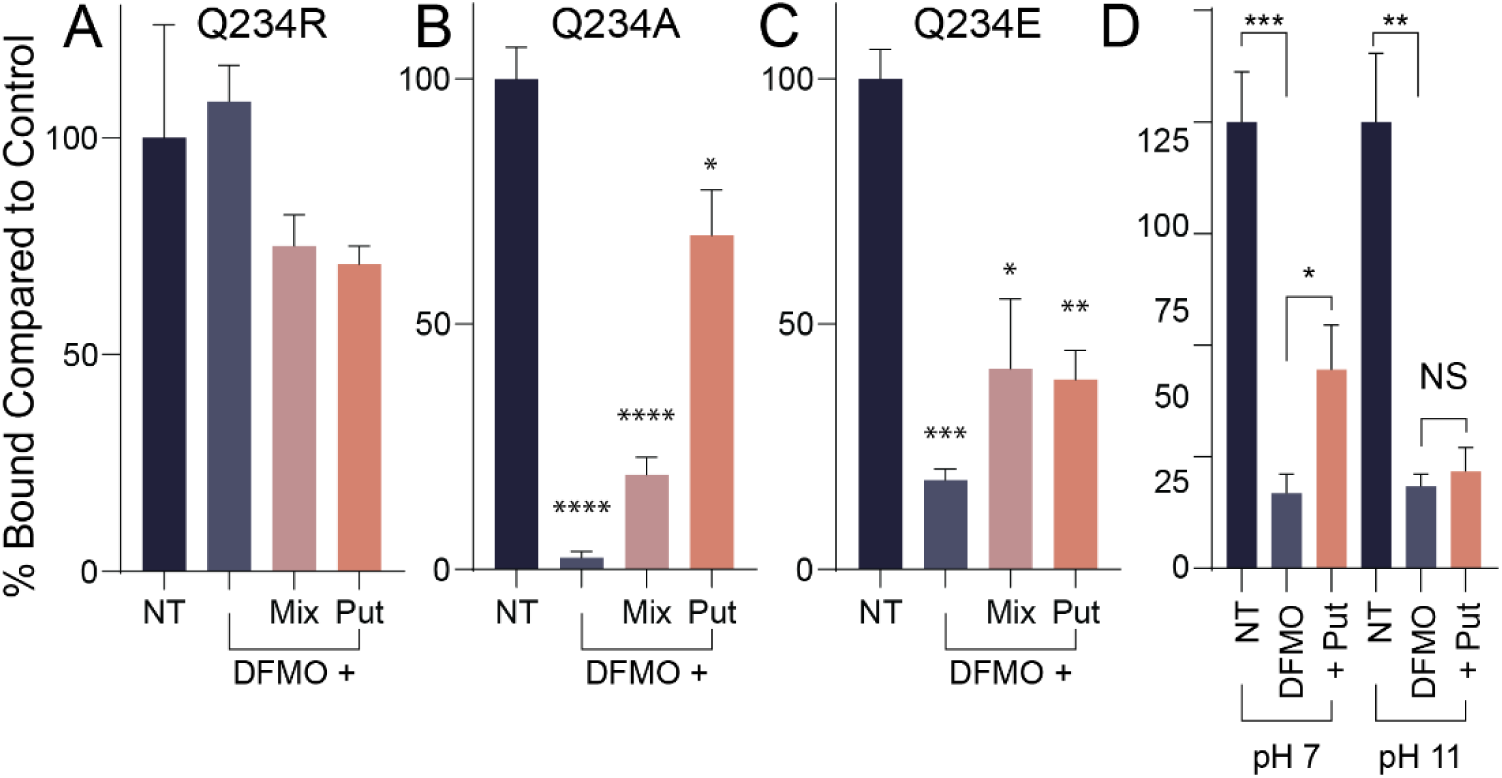
Positively-charged putrescine or Q234R mutation of VP3 mediates attachment. (A) Vero cells were treated with 1 mM DFMO for four days prior to attachment assay with CVB3 incubated with or without 5 mM putrescine at pH 7 or pH11. Attached virus was revealed and quantified after plaque formation. (B) Cells were treated as in (A) and subsequently infected with infectious clone-derived VP3^Q234R^, VP3^Q234A^, or VP3^Q234E^ CVB3. Attachment was measured as in (A). (D) Attachment assay was performed as in (A) but inoculum was incubated at pH 7 or pH 11 during incubation and attachment. NS not significant, *p<0.05, **p<0.01, ****p<0.0001 by Student’s T test (N≥2).

**Figure 5.**
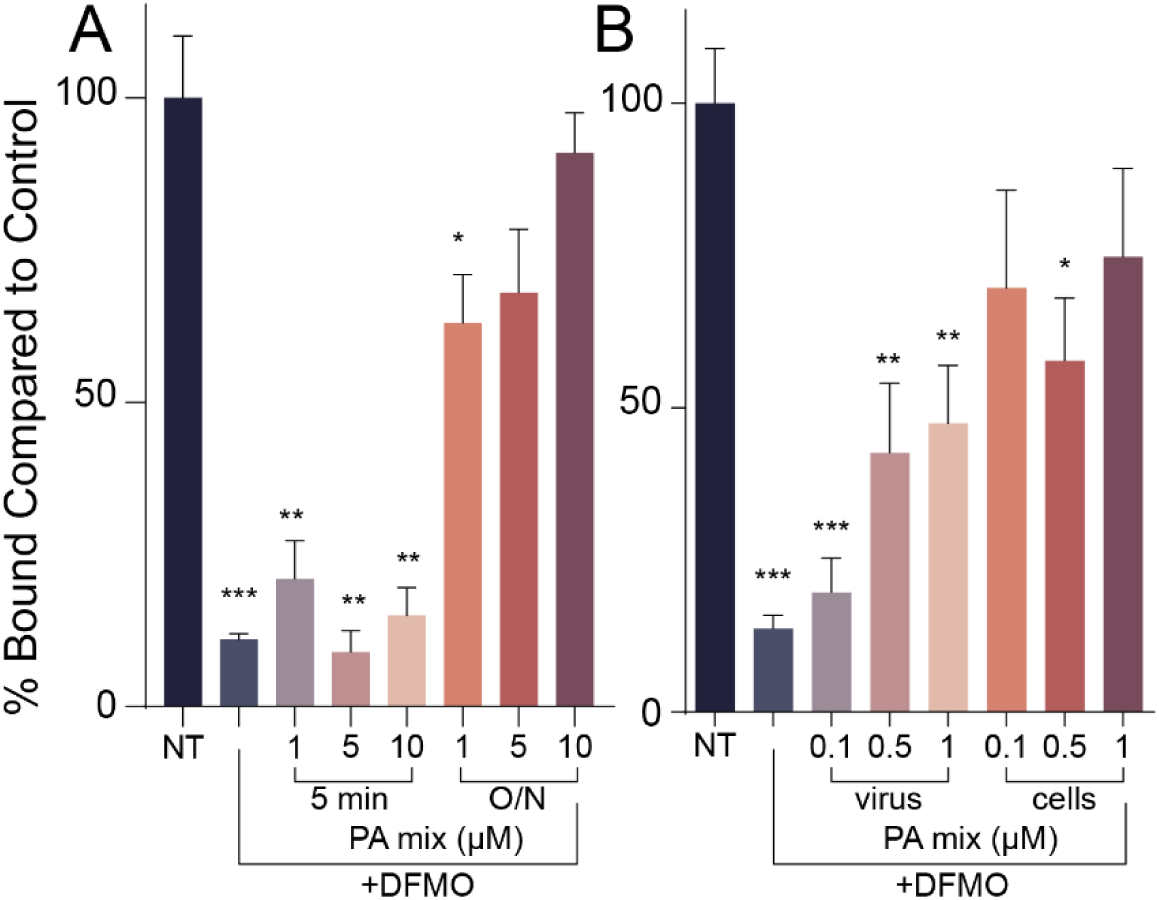
Cellular factors contribute to polyamine-mediated attachment. Vero cells were treated with 1 mM DFMO for four days prior to measuring CVB3 attachment. (A) Cells were incubated with exogenous polyamines as a mixture of spermidine, spermine, and spermine for either 5 min or overnight (O/N). (B) cells were treated as in (A) but either virus or cells were treated with a mixture of polyamines immediately prior to attachment assay. *p<0.05, **p<0.01, ***p<0.001, ****p<0.0001 by Student’s T test (N≥3).

### Putrescine’s positive charge mediates viral attachment

At physiological pH, polyamines are positively charged. We considered that the positive charge on putrescine likely mediates viral attachment and that neutralizing this charge would abrogate viral attachment. To test this, we incubated virus and putrescine together as previously, but we increased the pH to 11 to reduce the amount of positive charge on putrescine. We also incubated virus without putrescine at pH 11 to control for any effect of this change in pH on virus attachment itself. When we applied this virus to cells, we observed that in virus incubated with putrescine at pH 7, the molecule mediated attachment. However, incubation of CVB3 with putrescine at pH 11 eliminated the ability of the polyamine to mediate attachment (Figure 4D), suggesting that the positive charge on putrescine is required for it to facilitate CVB3 attachment.

### Cellular factors contribute to polyamine-mediated attachment

When we incubate virus with polyamines, we observe a rescue in virus attachment, specifically with putrescine and in a dose-dependent manner. We wished to investigate whether incubation of cells with polyamines directly might also rescue virus attachment, and to this end, we applied polyamine-containing media to DFMO-treated cells, removed this media, and then applied CVB3 for five minutes on ice, washing away unbound virus and adding agarose-containing overlay medium. Despite adding polyamines to the inoculum, we observed no rescue in virus attachment, suggesting that polyamines (including putrescine) may not mediate viral attachment when applied to cells. However, we previously demonstrated that adding polyamines to the cellular media 16h prior to viral attachment rescued virus replication. We recapitulated these results, which suggest that polyamine rescue of viral attachment requires a prolonged incubation time, perhaps suggesting that cellular processes dependent on polyamines need to modulated to mediate viral attachment.

### Polyamines facilitate heparan sulfate cell-surface presentation

Our prior work showed that diverse viruses rely on polyamines for attachment, including Zika virus, human rhinovirus, and Rift Valley fever virus^20^. Given the diversity of viruses exhibiting this phenotype, we considered that perhaps polyamines facilitate the expression of a common attachment factor for each of these viruses. With this consideration, we investigated whether heparan sulfates, negatively-charged cell surface molecules involved in virus attachment, were impacted by polyamine depletion. We measured cell surface expression of heparan sulfates via immunofluorescence (Figure 6A), using an antibody specific to heparan sulfates. To these cells, we also added sodium chlorate, a sulfation inhibitor, as well as DFMO to deplete polyamines. We observed significant cell surface staining in untreated cells, but in DFMO-treated cells, we found that heparan sulfates were significantly reduced, to a level similar to sodium chlorate treatment.

**Figure 6.**
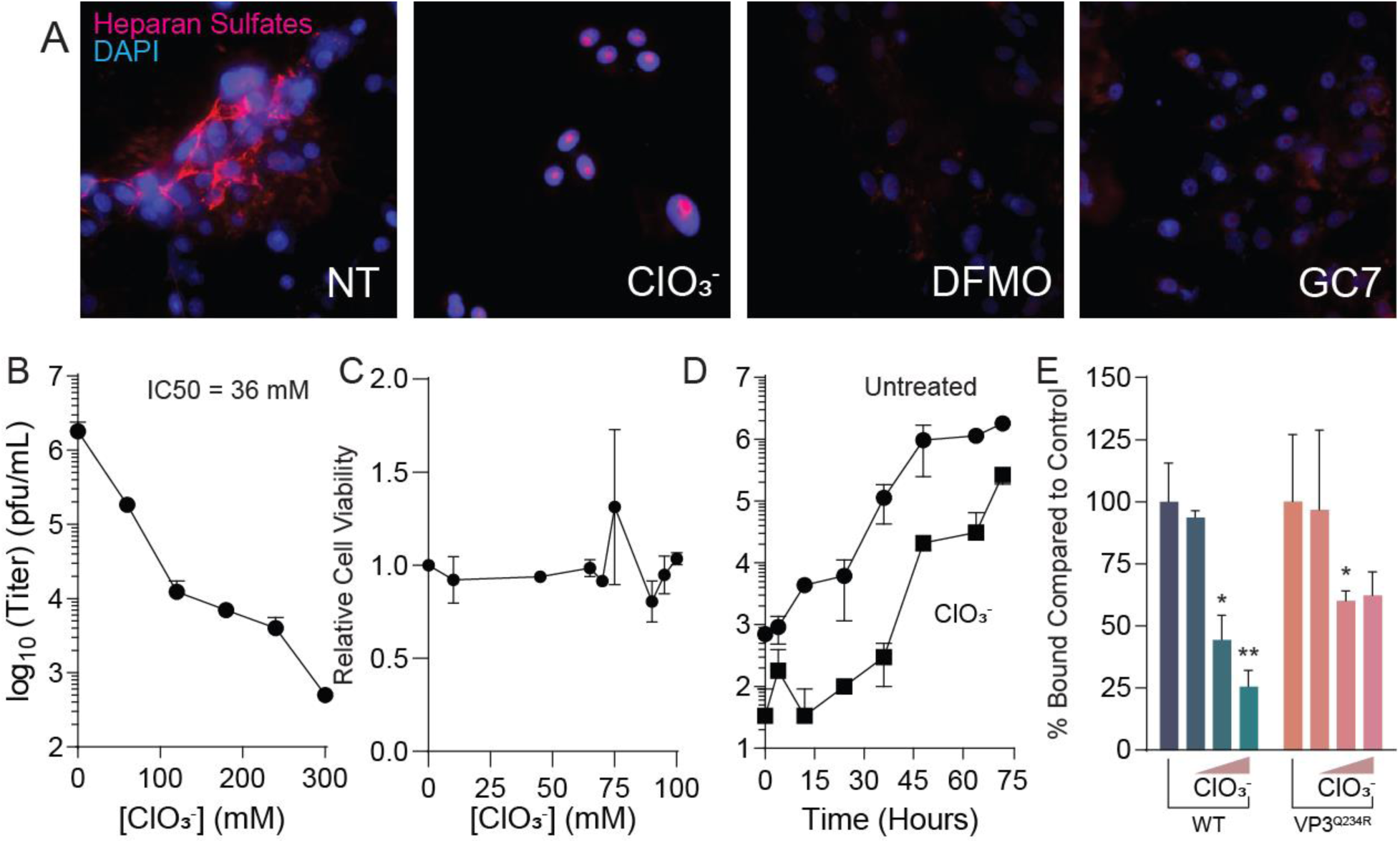
Polyamines facilitate cellular heparan sulfate synthesis. (A) Veros were treated with 100 mM chlorate (ClO3-), 1 mM DFMO, or 500 μM GC7 and subsequently stained for heparan sulfates and visualized by immunofluorescence. (B) Vero cells were treated with increasing doses of chlorate for four days prior to infection with CVB3 at MOI 0.1. Viral titers were determined at 48 hpi. (C) Cells were treated as in (A) and cellular viability measured. (D) Vero cells were treated with 100 mM chlorate and infected with CVB3 at MOI 0.1. Viral titers were determined at the times indicated. (E) Veros were treated with increasing doses of chlorate and WT and VP3^Q234R^ mutant CVB3 attachment was measured. *p<0.05, **p<0.01 by Student’s T test (N≥3).

**Figure 7.**
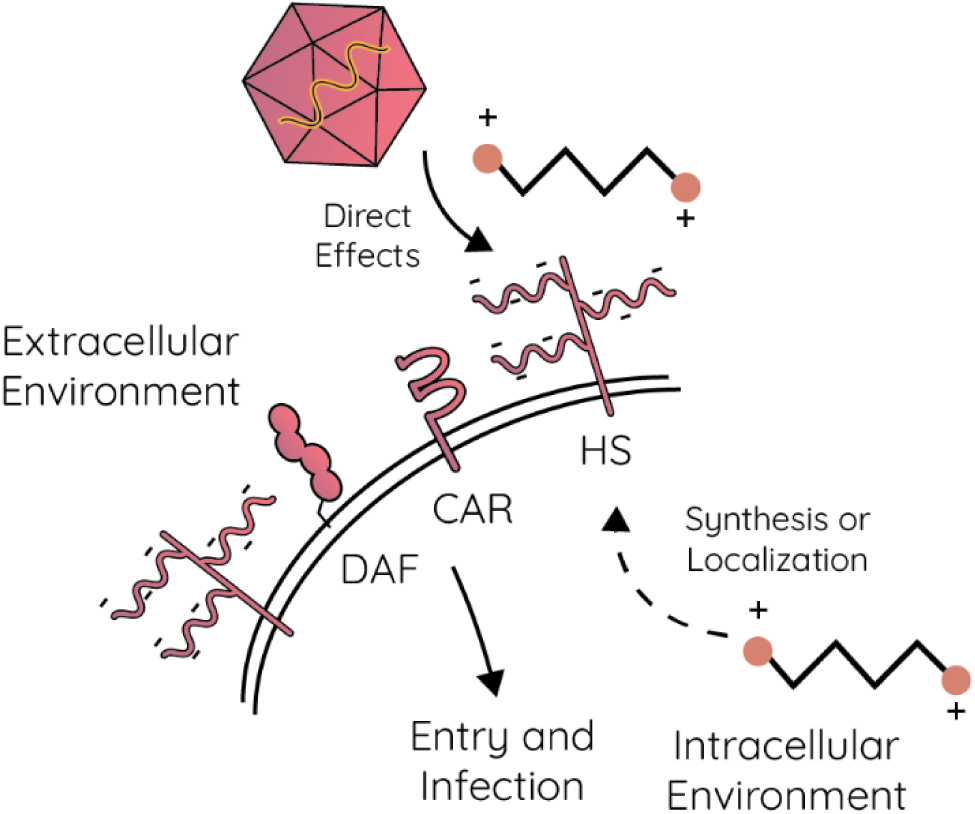
Working model. Polyamines facilitate CVB3 attachment and entry by directly mediating cellular attachment through the positive charge of the polyamines. Polyamines also support synthesis and localization of heparan sulfates, a nonspecific attachment factor, to facilitate CVB3 attachment.

To begin understanding how polyamines affect heparan sulfate synthesis and presentation, we measured expression of a variety of heparan sulfate synthetic genes, involved both in the synthesis of the core protein, polymerization of carbohydrate moieties, and sulfation of those carbohydrates. In all scenarios, we observed no significant change in gene expression in DFMO-treated cells, suggesting that perhaps DFMO does not regulate heparan sulfate synthesis at the level of gene expression. However, polyamines are involved in translation through the unique modification of eIF5A called hypusination, in which spermidine is conjugated to eIF5A and carboxylated. To determine if hypusination contributes to heparan sulfate synthesis, we imaged heparan sulfates by immunofluorescence using a specific inhibitor of hypusination called GC7. GC7 inhibits the first step in hypusination (conjugation of spermidine to eIF5A by deoxyhypusine synthase). Interestingly, we observed that heparan sulfates were significantly reduced in GC7-treated cells, suggesting that hypusination contributes to heparan sulfate synthesis and/or modification.

The Nancy strain of CVB3 has been reported to rely on heparan sulfates for attachment, but to confirm that heparan sulfates are important for virus attachment and infection, we first measured virus replication in sodium chlorate-treated cells. When treating cells with increasing doses of sodium chlorate, we observed a dose-dependent decrease in CVB3 titers (Figure 6B), with modest impacts on cellular viability (Figure 6C). This was true over several rounds of virus replication, as viral titers were reduced with sodium chlorate treatment over a timecourse (Figure 6D). Similarly, we found that sodium chlorate reduced virus attachment in a dose-dependent manner (Figure 6E). We used our VP3^Q234R^ mutant in these attachment assays and observed a modest rescue in viral attachment in these cells, suggesting that a portion of this mutant’s resistance may originate from its ability to bind cells depleted of heparan sulfates or that the CVB3 VP3^Q234R^ mutant has overall enhanced cellular attachment. In sum, these data highlight the role of polyamines and hypusination in heparan sulfate synthesis.

## Discussion

Our prior work with CVB3 showed that polyamines facilitate at least one step during viral attachment and entry, as depletion of polyamines limits the amount of virus associated with susceptible cells^20^. This phenotype is reverse with exogenous polyamines, though it was unclear whether polyamines directly or indirectly facilitate attachment. Our current work demonstrates that polyamines function both at the level of directly enhancing virus-cell association and by facilitating the synthesis of cellular heparan sulfates, a common attachment factor for several viruses^1,5^. Thus, these results likely hold true for other viruses that rely on heparan sulfates for attachment, as we observed previously.

We find that specifically putrescine, the “simplest” of the polyamines, facilitates direct attachment of virus to cells. Precisely why putescine, rather than the other polyamines with more charges and longer carbon chains, specifically enhances attachment is unclear, though one could hypothesize that this polyamine specifically associates with the charge landscape of the enterovirus virion. The capsid proteins exhibit canyons and valleys that mediate interaction with cellular molecules, including heparan sulfates and the cellular receptors. Whether polyamines and specifically putrescine bind to a specific portion of the CVB3 capsid is unclear. Structural analysis of the viral capsid proteins has never cocrystalized a polyamine with a structural protein; however, these proteins are often highly purified and may lose polyamine association upon purification.

Passaging CVB3 in polyamine-depleted cells, treated with DFMO, we observed a mutation in VP3 that confers resistance through enhanced cellular attachment^20^. The VP3^Q234R^ mutation was previously described to enhance association with the CVB3 receptor CAR (Coxsackie and adenovirus receptor)^25,26^, which we hypothesized was overcoming a deficit in cells treated with DFMO. It is tantalizing to consider that polyamines, specifically putrescine, may facilitate VP3’s association with CAR, and that upon polyamine depletion, CVB3 adapts by conferring a positive charge to this amino acid, negating the requirement for putrescine. Current work is addressing this model.

The connection between polyamines and other metabolic pathways has been established by our^27^ and others’ research^28,29^. Polyamines have previously been shown to associate with heparan sulfates and that cells can take up polyamines from the extracellular environment via heparan sulfates^30–33^. We find that heparan sulfate synthesis and surface expression is facilitated by polyamines, as treatment of cells with DFMO diminishes their presence on cells. When we investigated whether the expression of heparan sulfate synthesis or modification genes, we found no significant changes in expression (not shown); however, the regulation of heparan sulfate synthesis could be at the level of translation, through hypusination of eIF5A^34^. Whether polyamines facilitate the expression and presentation of other viral attachment factors on the surfaces of cells remains incompletely understood, but future work will need to characterize cell surface molecules impacted by polyamines and how this impacts virus attachment.

Several molecules like polybrene or protamine sulfates enhance viral attachment to cells and are frequently used to enhance transduction efficiency^35–37^. Interestingly, the structure of polybrene is highly similar to polyamines, comprised of carbon chains with quaternary amine groups, which confers a repeating positive charge. Polybrene has been shown to enhance attachment of viruses to cells in the absence of viral receptors^35^. Additionally, polybrene may act by aggregating viruses, enhancing their potential to productively infect^38^, or by neutralizing the negative cell surface charge (potentially through negatively-charged heparan sulfates)^39^ to enhance virus association. Thus, polyamines and putrescine in particular may be functioning similarly. In our assays, we used high levels of polyamines, in the millimolar range^40,41^; however, cellular levels of polyamines can be in this range. As a pathogen transmitted by the fecal-oral route, CVB3 also likely encounters high polyamine levels within the intestinal tract, where both bacterial and mammalian cells produce polyamines^42–45^. While our work does not address the *in vivo* physiological relevance of polyamines during natural infection with CVB3, it highlights a novel role for polyamines in both directly and indirectly facilitating viral attachment.

## Acknowledgments

This work was supported by R35GM138199 from NIGMS (BCM). We thank Dr. Ivana Kuo for assistance with microscopy.

## Materials and Methods

### Cell culture

Cells were maintained at 37°C in 5% CO_2_, in Dulbecco’s modified Eagle’s medium (DMEM; Life Technologies) with bovine serum and penicillin-streptomycin. Vero cells were obtained through Biodefense and Emerging Infections (BEI) Research Resources, NIAID, NIH (NR-10385), and were supplemented with 10% new-born calf serum (NBCS; Thermo Fisher).

### Drug treatments

Difluoromethylornithine (DFMO; TargetMol) and N^1^,N^11^-Diethylnorspermine (DENSpm; Santa Cruz Biotechnology, Santa Cruz, CA, USA), GC7 and sodium chlorate (Sigma-Aldrich, St. Louis, MO) were diluted to 100× solution (100mM, 10mM, 50 mM, and 10 M, respectively) in sterile water. For DFMO treatments, cells were trypsinized (Zymo Research) and reseeded with fresh medium supplemented with 2% NBCS. Cells were treated with 1 mM DFMO unless otherwise indicated. Cells were incubated with DFMO for 96 h, DEMSpm for 16 h, GC7 for 16 h, or sodium chlorate for 96 h to allow for depletion of polyamines or heparan sulfates. Experiments involving polyamine rescues were performed using 10 µM polyamines (Sigma-Aldrich) unless otherwise indicated and added to either the cell supernatant or viral inoculum, as indicated. The polyamine mix is a 1:1:1 equimolar solution of putrescine, spermidine, and spermine.

### Infection and enumeration of viral titers

CVB3 (Nancy strain) was derived from the first passage of virus in Vero cells after rescue from an infectious clone. VP3^234R^, VP3^234E^, and VP3^234A^ mutants were generated as previously described^20^ using the following primers: VP3^234A^, 5’-CCT-TTC-ATT-TCG-GCC-AAC-TTT-TTC-C-3’ (F) and 5’-CCC-TGG-AAA-AAG-TTG-GCC-TGC-GAA-ATG-3’ (R); VP3^234E^, 5’-CCT-TTC-ATT-TCG-CAG-GAA-AAC-TTT-TTC-C-3’ (F) and 5’-CCC-TGG-AAA-AAG-TTT-TCC-TGC-GAA-ATG-3’ (R). For all infections, DFMO, DENSpm, sodium chlorate, and GC7 were maintained throughout infection as designated. Viral stocks were maintained at -80°C. For infection, virus was diluted in serum-free DMEM for a multiplicity of infection (MOI) of 0.01 on Vero cells, unless otherwise indicated. Viral inoculum was added to the cells and supernatants were collected at specified time points. To quantify viral titers via plaque assay, dilutions of cell supernatant were prepared in serum-free DMEM and used to inoculate confluent monolayers of Vero cells for 10 to 15 min at 37°C. Cells were overlain with 0.8% agarose in DMEM containing 2% NBCS. CVB3, VP3^234A^, VP3^234E^ samples were incubated for 2 days and the VP3^234R^ mutant for 3 days at 37°C. Cells were fixed with 4% formalin and revealed with crystal violet solution (10% crystal violet; Sigma-Aldrich). Plaques were enumerated and used to back calculate the number of PFU per ml of collected volume.

### Plaque formation attachment assay

Vero cells were seeded in 6-well plates and grown to confluence in DMEM with 2% NBCS. The cells were treated for 96 h with 1 mM DFMO. For polyamine rescue experiments, cells were treated overnight before the infection with 10 µM or incubated with the viral inoculum for 5 min at room temperature with .1, .5, 1, or 5mM of polyamines (Millipore Sigma) unless otherwise indicated. After the 96-h DFMO treatment, the cells were placed on ice and the medium was aspirated from the cells and replaced with .5 ml serum-free medium containing either 1,000 or 2,000 PFU. The infected cells were incubated on ice for a specified amount of time. After the specified time, the cells were washed 3x with PBS and then overlaid with 0.8% agarose containing DMEM with 2% NBCS. The plates were incubated at 37°C for plaques to develop. CVB3, VP3^234A^, and VP3^234E^ were incubated for 2 days; the VP3^234R^ mutant for 4 days. The cells were fixed with 4% formalin, and the plaques were visualized with crystal violet staining.

### qPCR based attachment assay

Vero cells were seeded at 1.5 × 10^5^ cells per well in 12-well plates in DMEM with 2% NBCS. The cells were treated for 96 h with 1mM DFMO or 16 h with DENSpm. After 96 h or 16 h, the media were aspirated from the cells and replaced with 100 μL of serum free media containing virus. The infected cells were incubated for 10 min at room temperature or on ice. The cells were then washed 1× with PBS, and then, 200 μL of Trizol was added to the cells. The RNA was extracted with the Zymo RNA extraction kit, converted to cDNA, and quantified by real-time PCR with SYBR Green (DotScientific) using the one-step protocol QuantStudio 3 (ThermoFisher Scientific). Relative genomes were calculated using the ΔCT method, normalized to the β-actin qRT-PCR control, and calculated as the fraction of the unwashed samples. Primer sequences are as follows: CVB3, 5’-AGG-GCG-AGA-TCA-ATC-ACA-TTA-G-3’ (F) and 5′-CTC-TGC-TGT-TGC-CTC-ACT-ATC-3′ (R); β-actin, 5′-CAC-TCT-TCC-AGCCTT-CCT-TC-3′ (F) and 5′-GTA-CAG-GTC-TTT-GCGGAT-GT-3′ (R). Primers were verified for linearity using 8-fold serial diluted cDNA and checked for specificity via melt curve analysis.

### Immunofluorescence imaging

Cells grown on coverslips were either treated with 1 mM DFMO, 100mM sodium chlorate, 10 µM DENSpm, 500µM GC7 or untreated. Cells were fixed with 4% formalin for 15 min, washed with PBS, permeabilized, and blocked with 0.2% Triton X-100 and 2% BSA in PBS (blocking solution) for 30 min at room temperature (RT). Cells were sequentially incubated as follows: primary mouse anti-10E4 (Amsbio, 370255-S) with blocking for 2 h at room temperature. Cells were subsequently washed then incubated with secondary goat anti-mouse antibodies (1:500 with blocking, 30 min, RT). Mounting media with DAPI was used to visualize nuclei. Samples were imaged with a Zeiss Axio Observer 7 with Lumencor Spectra X LED light system and a Hamamatsu Flash 4 camera using appropriate filters using Zen Blue software with a 40× objective.

